# Selection on early survival does not explain germination rate clines in *Mimulus cardinalis* (Phrymaceae)

**DOI:** 10.1101/2021.12.14.472651

**Authors:** Christopher D. Muir, Courtney L. Van Den Elzen, Amy L. Angert

## Abstract

**Premise:** Many traits covary with environmental gradients to form phenotypic clines. While local adaptation to the environment can generate phenotypic clines, other nonadaptive processes may also. If local adaptation causes phenotypic clines, then the direction of genotypic selection on traits should shift from one end of the cline to the other. Traditionally genotypic selection on non-Gaussian traits like germination rate have been hampered because it is challenging to measure their genetic variance.

**Methods:** Here we used quantitative genetics and reciprocal transplants to test whether a previously discovered cline in germination rate showed additional signatures of adaptation in the scarlet monkeyflower (*Mimulus cardinalis*). We measured genotypic and population level covariation between germination rate and early survival, a component of fitness. We developed a novel discrete log-normal model to estimate genetic variance in germination rate.

**Results:** Contrary to our adaptive hypothesis, we found no evidence that genetic variation in germination rate contributed to variation in early survival. Across populations, southern populations in both gardens germinated earlier and survived more.

**Conclusions:** Southern populations have higher early survival but this is not caused by faster germination. This pattern is consistent with nonadaptive forces driving the phenotypic cline in germination rate, but future work will need to assess whether there is selection at other life stages. This statistical framework should help expand quantitative genetic analyses for other waiting-time traits.

## Introduction

Populations within species differ in ecologically important traits that often result from adaptation to different local environments (Turesson, 1922; Clausen et al., 1948). However, it is still rare that we understand the traits and selective agents underlying local adaptation (Wadgymar, Lowry, et al., 2017). The classic signature of local adaptation is crossing reaction norms for fitness measured in a reciprocal transplant experiment (Kawecki and Ebert, 2004; Johnson et al., 2021), such that populations have higher relative fitness in their local environment but lower relative fitness in a foreign environment. For example, annual inland ecotypes of the yellow Monkeyflower *Mimulus guttatus* have higher relative fitness in seasonally dry sites compared to perennial coastal ecotypes that are adapted to year-round water availability. The relative fitness of coastal and inland ecotypes is reversed on the coast because of life history and other genetic differences between ecotypes (Lowry and Willis, 2010). The prevalence of local adaptation implies that selection varies spatially, populations have heritable variation in fitness, and selection is stronger than migration or drift. Exceptional cases of local adaptation over short temporal or spatial scales (e.g. Wright et al., 2013; Grant and Grant, 2014; Richardson et al., 2014; Lescak et al., 2015; Barrett et al., 2019; DiVittorio et al., 2020) likely contribute to an availability bias that leads many to perceive that local adaptation is ubiquitous. However, systematic meta-analyses reveal that local adaptation is often weak or nonexistent (Leimu and Fischer, 2008; Hereford, 2009; Brady et al., 2019) and may be getting weaker because of anthropogenic climate change (Bontrager et al., 2020). Populations may not be locally adapted if differential selection is weak relative to migration or drift or they lack heritable variation in traits under selection. Alternatively, recent anthropogenic climate change may have erased the signature of local adaptation by causing some foreign populations to have higher fitness than local populations (Bontrager et al., 2020). For example, in the alpine plant *Boechera strica*, simulated climate change (mimicking early snow melt) favors populations from lower elevations and snow addition restores the relative fitness advantage of high-elevation populations (Anderson and Wadgymar, 2020). Understanding what causes trait variation among populations within species, whether adaptive or not, will help biologists better predict how populations will respond to environmental change.

Phenotypic clines are commonly interpreted as evidence for local adaptation, but nonadaptive processes also generate similar patterns (Endler, 1977). A phenotypic cline is a correlation between a heritable character and the environment or a proxy for environmental gradients such as latitude or elevation (Huxley, 1938). Clines may be caused by either abrupt or gradual environmental change. For example, heavy metal contamination of soil around mines is an abrupt change from the surrounding pasture soil. A similarly abrupt change in the heavy metal tolerance of *Anthoxanthum odoratum* occurs across the mine boundary, but other traits under weaker or correlated selection vary gradually over space (Antonovics and Bradshaw, 1970). Other clines track gradual environmental change. Latitudinal clines in the size of *Drosophila subobscura* flies have evolved multiple times and most likely track gradual variation in temperature and/or phenology (Huey et al., 2000). But clines are not necessarily adaptive. When there is genetic isolation by distance (Wright, 1943), it is possible for phenotypic differences to correlate with genetic differences throughout the species range (Vasemägi, 2006). In practice, nonadaptive clines are difficult to demonstrate because it is hard to reject adaptive explanations. The proportion of cyanogenesis varies clinally in white clover (*Trifolium repens*) but field experiments have not observed spatially varying selection that could explain this cline (Wright et al., 2021). Hence, spatial variation in cyanogenesis may be nonadaptive or the design of the experiment may have missed selection occurring during germination and early development. When species expand their range, such as during biological invasions, nonadaptive clines can readily evolve because of multiple introductions and serial founder events (Colautti and Lau, 2015). Even parallel clines, which are usually considered strong evidence of natural selection, can result from nonadaptive processes when there is epistasis and consistent spatial variation in the strength of genetic drift (Santangelo et al., 2018). Evolutionary ecological experiments such as reciprocal transplants are needed to provide additional evidence to distinguish between adaptive and nonadaptive hypotheses for clines (Wadgymar, Daws, et al., 2017).

There are several experimental methods for measuring the agents and strength of selection to different environments under natural conditions (Wadgymar, Lowry, et al., 2017), but here we focus on differential selection. One key prediction is that the direction of selection and position of the phenotypic optimum will differ between environments. If genetic variation in a trait *causes* fitness to vary, then genotypic selection analyses (Rausher, 1992) should find that genotypes with high trait expression are favored in one environment but disfavored in another. For example, selection favors higher seed dormancy in southern environments of *Arabidopsis thaliana* and lower dormancy in northern environments, presumably to optimally time seedling establishment with the onset of favorable summer conditions (Postma and Ågren, 2016). Populations can also simultaneously differ as a result of both adaptive and non-adaptive processes. When locally adapted populations differ in traits that do not themselves confer local adaptation, we expect population-level covariation between divergent traits and fitness but do not expect within-population variation to correlate with fitness. Hence, comparing the direction of among-genotype and among-population selection can help distinguish between phenotypic clines that are important for local adaptation and those that may have evolved nonadaptively.

In this study, we used a reciprocal transplant to test whether a cline in *Mimulus cardinalis* germination rate could be locally adaptive in terms of early survival between seedling establishment and flowering. Because seeds of *M. cardinalis* do not require cold or warm stratification to break dormancy, they will germinate whenever conditions are sufficiently warm and moist. Within this germination niche, there is genetic variation for germination time that covaries with latitude and other traits associated with a fast-slow growth continuum (Muir and Angert, 2017; Sheth and Angert, 2018; Nelson et al., 2021). In greenhouse conditions, seeds from the southern end of the range (San Diego County, CA) germinate in 6–7 days; seeds from the northern end of the range (Lane County, OR) germinate in 9–10 days. The growing season is typically longer in the southern end of the range because of reduced or nonexistent snowpack, but for the same reason southern populations may be more susceptible to late-season drought (Sheth and Angert, 2018). The adaptive hypothesis is that faster germination increases early survival in the southern end of the range and slower germination increases early survival at the northern end. We would predict either stabilizing genotypic selection for different optimal germination rates or opposing signs of directional genotypic selection. If the cline is maintained by nonadaptive forces then germination rate should be uncorrelated with genotypic fitness (even if it is correlated with population mean fitness) or the direction of selection should be the same in both parts of the range. A limitation of this study is that we cannot measure selection on seedling establishment since that occurred in the greenhouse (see Materials and Methods) or later survival and fecundity (these fitness components will be analyzed in a follow-up paper). Germination and early survival are potentially important for local adaptation but often overlooked compared to later life stages. Failure to measure seedling establishment and early survival can bias estimates of selection on traits that only express at later life stages, such as flower size (Mojica and Kelly, 2010). In the herb *Arabidopsis thaliana*, quantitative differences in germination timing contributed to local adaptation and are maintained by a balance of selection on seedling establishment, survival, and fecundity, but the optimal timing varies in northern and southern environments (Postma and Ågren, 2016, 2018). A challenge of measuring genotypic selection on germination rate is that quantitative genetic theory and analysis are built around Gaussian distributions (Lynch and Walsh, 1998; but see Villemereuil et al., 2016), but the distribution of times until germination are most likely not Gaussian. Although it is convenient to express germination as a rate (events per time), this and other “waiting time” traits are measured as the time until an event, the inverse of germination rate. Throughout this paper we distinguish between germination rate as the trait of interest which we estimate by measuring the time to germination. The time until germination (or another event) can be modeled as a waiting time, which is bounded at zero and usually right skewed. Waiting times are distributed exponentially if the probability of germination is constant through time, but this unlikely to be true. Rather, the probability of germination almost certainly increases through time once conditions are conducive. Hence, more complex waiting time distributions should occur in nature. A second challenge is that waiting time is usually modeled as a continuous trait, but is observed at discrete intervals (e.g. once per day). If the average time to germination is sufficiently long relative to the interval between observations, then a continuous-time approximation may be sufficient. However, when time to germination is several days, as with *M. cardinalis*, then accurately modeling the trait may need to account for the discrete observation process. Here we show that non-Gaussian distributions and measurement process can be incorporated into and improve analysis of time to germination. This approach should be useful, with some customization, to many other traits and biological systems. Hence, investigating spatially variable selection on germination time is important for expanding our knowledge of local adaptation and phenotypic clines for poorly studied life stages with challenging statistical properties.

## Materials and Methods

### Study system

*Mimulus cardinalis* (=*Erythranthe cardinalis* Lowry et al., 2019) is a perennial forb native to the Western US (California and Oregon). It is self-compatible, predominantly outcrossing, and hummingbird pollinated. We used five focal populations from throughout the geographic range of *M. cardinalis* (Table 1; Fig. S1). These five focal populations are a subset of those we used previously to identify a cline in germination and other traits (Muir and Angert, 2017). Seeds were collected in the field from mature, undehisced fruits left in open coin envelopes for 2 − 4 weeks to dry, then stored at room temperature. To minimize maternal effects, we grew a large number of field-collected seeds in the greenhouse and generated seed families for this experiment by hand-pollinating individuals using the breeding design described in the next section.

**Table 1:** Source populations, including the name of the drainage where the seeds were collected, the latitude, longitude, and elevation in meters above sea level (mas).

| Source population | Latitude | Longitude | Elevation (mas) |
| --- | --- | --- | --- |
| Sweetwater River | 32.900 | −116.585 | 1180 |
| West Fork Mojave River | 34.284 | −117.378 | 1120 |
| North Fork Middle Tule River | 36.201 | −118.759 | 926 |
| Little Jamison Creek | 39.743 | −120.704 | 1603 |
| Rock Creek | 43.374 | −122.957 | 326 |

### Genetic variance and heritability

To estimate selection among genotypes and populations, we first need to quantify genetic variance in germination rate and early survival. We estimated genetic variance and heritability of germination rate and winter survival using a quantitative genetic breeding design. For each population, we crossed 15 parental individuals using a partial diallel design (Lynch and Walsh, 1998) with three dams per sire and three sires per dam for a total of 45 full-sib families per population. The number of parents is low for quantitative genetic analyses, but we were restricted garden space and by the availability of wild-collected seeds in some source populations. We opted for fewer parental individuals and more populations so that we can better understand range-wide patterns of local adaptation. Individual plants of this hermaphroditic species were used as both sires (pollen parent) and dams (ovule parent). We did not make crosses between populations. One family from the Little Jamison Creek population did not produce enough seeds, resulting in 44 usable families. In total, we used 5 × 45 − 1 = 224 families.

We estimated genetic variances and heritabilities for each of the five populations (Table 1). The total phenotypic variance within a population *V*_*p*_ can be partitioned into genetic (*V*_*G*_), maternal (*V*_*M*_), and environmental (*V*_*E*_) components:

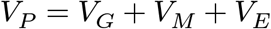

Genetic and environmental factors contribute to maternal affects (Lynch and Walsh, 1998). Although we attempted to minimize maternal effects by growing a refresher generation in a common greenhouse environment and *M. cardinalis* seeds are tiny and not well provisioned, seed traits may be particularly sensitive to maternal effects. We did not have statistical power to estimate nonadditive genetic contributions, therefore we estimated broad-sense (*H*^2^) heritability, the fraction of phenotypic variance contributed by all sources of (including dominance and epistasis) genetic variance:

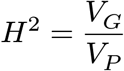

When separate individuals are used as dams and sires, additive genetic variance of the base population is often estimated from the variance among sires: *V*_*G*_ = 4 *σ*_sire_ (Lynch and Walsh, 1998). Since our breeding design used hermaphroditic individuals as both dams and sires, we estimated parental breeding values from their contributions as both sires and dams. Hence, we estimated *V*_*G*_ from the variance among parents:

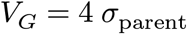

Using *σ*_parent_ rather than *σ*_sire_ is more powerful because it uses all of the data. We estimated *σ*_parent_ and other parameters using a Bayesian mixed effects model. We describe the general approach here and provide detail specific to germination rate and survival below. We fit the model in *Stan* version 2.29.2 (Stan Development Team, 2022) using **cmdstanr** version 0.5.2 (Gabry and Češnovar, 2022). *Stan* calculates the posterior distribution of the model using the Hamiltonian Monte Carlo algorithm, which is similar to the more widely used Markov Chain Monte Carlo, but is faster and more efficient at sampling for many applications (Monnahan et al., 2017). We used weakly informative normally distributed priors for parameters that affect the trait mean (intercepts and coefficients) and half-Cauchy priors for variance parameters. Weakly informative priors are strongly recommended for complex mixed models (McElreath, 2016). We ran 4 parallel chains for 4000 warmup iterations, 4000 sampling iterations, and a thinning interval of 4. This configuration allowed parameters to converge, which we defined as the convergence diagnostic 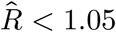 (Vehtari et al., 2021). We inspected posterior predictive plots to assess model adequacy using the pp_check function from the *R* package **bayesplot** version 1.9.0 (Gabry et al., 2019). We used the posterior median for point estimates and calculated uncertainty with the 95% quantile intervals from the posterior distribution. *Stan* and *R* code are available on GitHub (https://github.com/cdmuir/mimulus-germination) and will be archived on Zenodo upon publication. Data will be deposited on Dryad upon publication.

### Germination rate

We estimated germination rate under benign greenhouse conditions by measuring time to germination, the number of days to visible emergence from the soil. There were 48 randomized blocks split evenly between two time cohorts (North cohort: August 3–4 and South cohort: August 22–24, 2015) in the University of British Columbia greenhouse (Vancouver, BC, Canada). The North seed cohort went into the North transplant garden (gardens described below) and the South seed cohort went into the South transplant garden. Seed cohorts were staggered so that seedlings were transplanted at the same ontogenetic stage, as described below. We distinguish between cohorts and gardens because germination for each transplant garden was assayed in the greenhouse prior to transplanting. Emergence is a reliable proxy for germination since seeds were sown directly on top of the soil, resulting in little delay between germination and visible cotyledons. We sowed 3–5 seeds per plug on moist perlite-peat moss potting medium (Sunshine #1, Sungro Horticulture, Agawam, Massachusetts, USA) and recorded the first day on which a plug had at least one germinated seedling. We later thinned each plug to one seedling by selecting the central-most individual, so the first germinated seedling recorded is generally not the same individual transplanted into the field. Hence, individual correlations between germination and survival are inappropriate and we analyze relationships between breeding values or population averages. We censused germination daily for most of the experiment, but census dates were more spread out later in the experiment when only a few plants remained. We accounted for census interval in our statistical model (see below). Gentle misting kept soil moist for the duration of the experiment. Most plugs (9320 of 10650, 87.5%) had at least one germinant by the end of the experiment (North cohort: August 27, 2015; South cohort: September 20, 2015); we treated plants that did not germinate as missing data. 5 (0.05%) plants emerged before day 5. We removed data on these individuals because they were likely not *M. cardinalis*, but rather contamination from another species’ seed in the potting medium. Species can be difficult to tell apart as the cotyledons first emerge, but in plugs where a single germinant emerged before day 5 it was always a contaminant species. It therefore seems most likely that all germinants before day 5 were contamination, but observations during thinning showed that contamination was rare and it unlikely to have any substantial impact on our estimate of germination timing.

### Estimating genetic variance in germination rate

In this section we describe the custom probability distribution we used to estimate quantitative genetic parameters of germination rate in this study. However, this approach could be extended to other ecologically important non-Gaussian traits censused at discrete temporal or spatial intervals. To modify the approach for other systems requires specifying an appropriate statistical distribution for the trait of interest and using integration to calculate the probability of an event occurring within a definite time or spatial interval. Probabilistic programming languages like *Stan* (Stan Development Team, 2022) have made it easier for biologists to specify statistical models most appropriate for their data.

The full germination-rate model included a fixed effect of population, fixed effect of cohort, a random effect of greenhouse block, a random maternal effect of dam, and a random effect of parent as either dam or sire to estimate genetic variance. Most quantitative traits are modeled with a Gaussian (i.e. normal) distribution, but this is inappropriate for germination which has a lower bound at 0 and is censused at discrete time points. A Gaussian distribution might be an adequate approximation if the time to germination was longer and more spread out so that the effects of a zero lower bound and discrete sampling interval were inconsequential. However, given the rapid germination rate and small spread in timing for *M. cardinalis*, we used a different approach. Instead, we modeled genetic variation in a continuously distributed latent rate parameter that is measured as a discrete number of days to germination. Time to germination can be modeled similar to other waiting time or survival processes. There are several common waiting-time distributions with a lower bound at 0, but we used the log-normal distribution because unlike some other common distributions (e.g. exponential), the mean and variance of log-normal distribution are separate parameters. Hence, we can model the variance components separate from differences in the mean germination rate. We discretized the distribution using the definite integral of the probability density to calculate a probability mass. Let the probability density with parameters 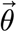 be 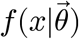. If a seed was censused to have germinated on day *t*_*i*_, it could have germinated at any time between the previous census on day *t*_*i*−1_. The associated probability mass *g*(*x*|*θ*) is:

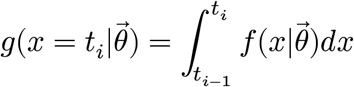

Preliminary analyses revealed a germination threshold of four days since we did not observe any germination before the fifth day after sowing. The model did not fit as well without this threshold because models without a threshold predict a significant amount of germination between days 0 and 5 (results not shown). The variances and heritability are reported on the log-transformed scale.

### Transplant gardens and winter survival

After germination, we transplanted 20 out of 24 blocks of seedlings each to two experimental gardens located at the northern and southern portions of the species’ range. We refer to these as the “North” and “South” gardens, respectively. We used some plants from the remaining 4 blocks to replace those that did not germinate or died during transport. We transplanted seedlings at similar ontogenetic stages but temporally staggered. We planted the North and South cohorts 5–6 weeks after germination on September 9–17, 2015 (North seed cohort into North garden) and October 2–9, 2015 (South seed cohort into South garden), respectively. We chose times when we observed natural seedlings were of similar size. The North garden was located near natural populations along the Middle Fork of the Feather River (39° 46’ 59.7” N, 120° 38’ 31.1” W, 1314 mas, Plumas County, California, USA); the South garden was located along King Creek (32° 54’ 23.4” N, 116° 37’ 26.0” W, 1011 mas, San Diego County, California, USA). These gardens are located close to two of the source populations (Fig. S1). The North garden is 7.1 km from the Little Jamison Creek source population; the South garden is 3.8 km from Sweetwater River source population. Since *M. cardinalis* is a riparian specialist, gardens were located within 50 m of natural waterways where the microclimate closely resembled that of nearby natural populations. We irrigated plots with microperforated drip tape (Toro, Bloomington, Minnesota) placed about each seedling to help establish transplants and mimic riparian soil where *M. cardinalis* naturally germinates. Seedlings were initially irrigated once per day. The duration of watering was adjusted at each garden to give each plant approximately the same volume of water. Once plants were established, we gradually reduced irrigation as evaporation declined while preventing drought stress. Irrigation ended when winter precipitation began. We discarded data on plants that died within the first month after transplanting, as this probably indicates transplant shock rather than natural mortality. We monitored reemergence in the spring using monthly censuses beginning on March 14, 2016 (South) and April 15, 2016 (North). If a plant was recorded in the last census of 2015 and emerged in spring 2016, we counted it as alive. If a plant did not reemerge, we recorded winter mortality.

### Estimating genetic variance in winter survival

We treated winter survival as a binomially distributed trait determined by a latent probability of survival *p*_surv_. We therefore estimated genetic variance in *p*_surv_ and then used methods described by Villemereuil et al. (2016) for non-Gaussian traits to calculate heritability. We analyzed Garden as a fixed effect and garden Block as a random effect. The primary difference from the germination model is that we estimated genotype-by-environment (G × E) interactions to test whether populations are locally adapted. If there is local adaptation, southern populations should have higher survival in the South garden and lower survival in the North garden, and *vice versa*.

### Selection on germination rate

We estimated genotypic and population-level selection on winter survival as a function of germination rate. Long-term demographic studies of natural *M. cardinalis* populations use size-rather age-based models, so we do not know with certainty how important a fitness component winter survival is. However, population growth rate is moderately sensitive to the survival of prereproductive individuals of uncertain age; early survival is less important than recruitment, but more important than fecundity (Angert, 2006). If genetic variation in germination rate within and between populations *causes* the probability of survival to change, then there should be significant genotypic selection on germination rate. Alternatively, populations with different mean germination rates may have different probabilities of survival as a consequence of other, confounding trait differences. If this is true, we predict a population-level correlation between germination rate and survival, but no evidence of genotypic selection on variation within populations. For each sample of breeding values and population averages from the posterior distribution, we used ordinary linear regression to calculate the relationship between germination rate and winter survival in both gardens at both genotypic and populational levels. We also calculated the variance among populations (*V*_pop_) in germination rate and survival from the posterior distribution. We used the median and quantile intervals to estimate the slopes and quantify uncertainty in our estimates. We used the same approach to estimate quadratic selection coefficients as well.

### Climate

We compared the climate during the experiment (September 2015 to May 2016) to a standardized climate normal. Following Sheth and Angert (2018), we downloaded climate variables for 1961–1990 and 2015–16 derived from ClimateNA version 7.10 (Wang et al., 2016). We compared the seasonal temperature average (autumn: September-November; winter: December-February; spring: March-May) for the experiment with the 1961–1990 normal for each population.

## Results

### Southern source populations germinate faster and have a higher probability of winter survival

The vast majority of seeds germinated in 1–2 weeks, but northern source populations took a few days longer on average. 4206 out of 5321 plugs (79.0%) had germinants in the North cohort; 5114 out of 5328 plugs (96.0%) had germinants in the South cohort. South cohort seedlings also germinated about one day faster (mean = 7.7, median = 7) than the North cohort (mean = 8.9, median = 8). The estimated log-mean difference and 95% CIs in time to germinate in the North cohort was 0.395 (0.284, 0.505). After accounting for cohort and block effects, southern source populations consistently germinate faster (Fig. 1a). For example, seeds from the slowest population (Rock Creek) germinated 2.9 (95% CI [1.8, 4.2]) days slower than the fastest population (West Fork Mojave River). The discretized log-normal distribution was an adequate model of germination rate based on the similarity of the posterior predictions to the observed data (Fig. S2).

**Figure 1:**
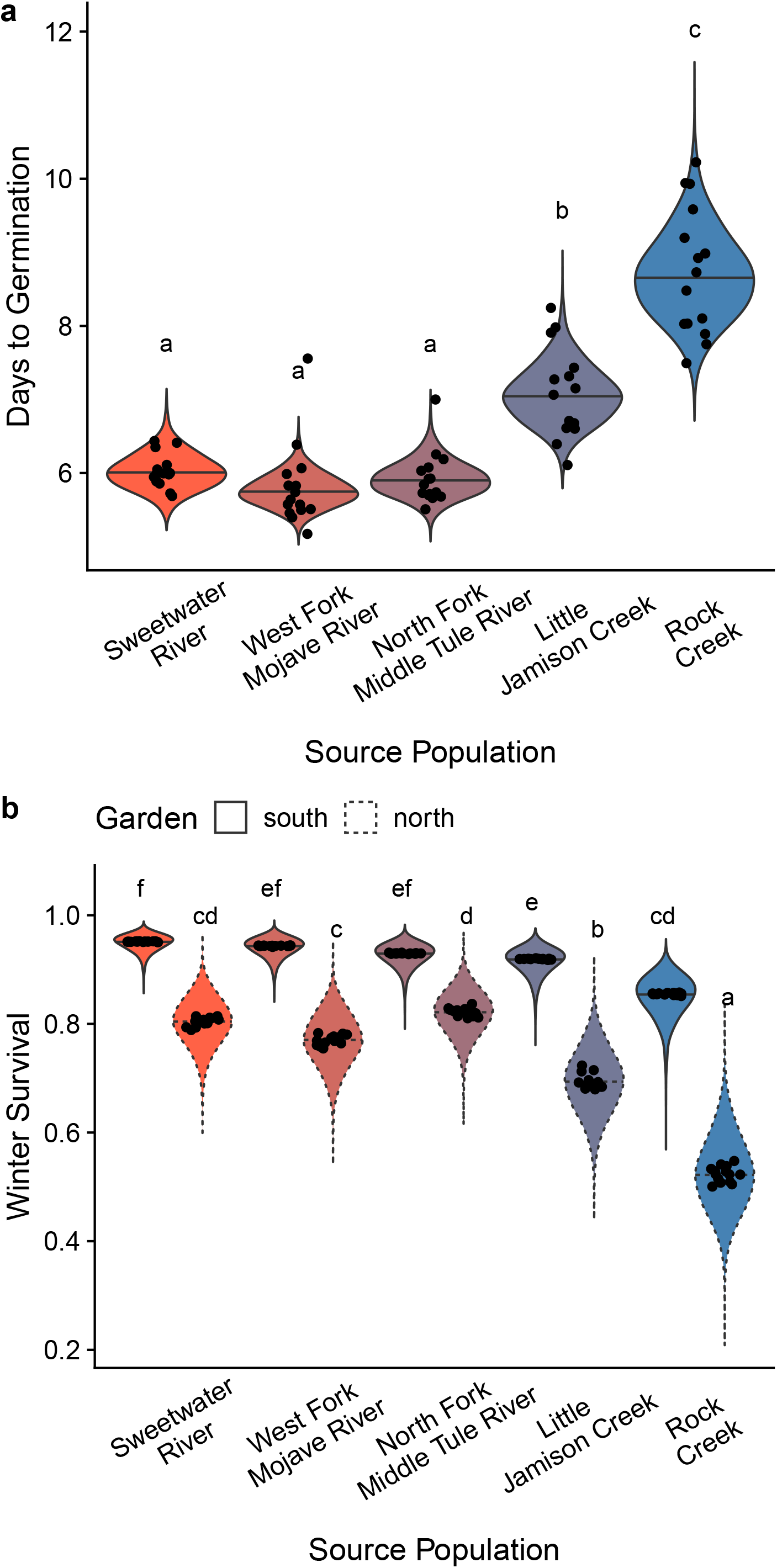
Germination rate and survival differ among source populations of *Mimulus cardinalis*. Source populations are arrayed along the *x*-axis by latitude of origin from south to north. In both panels, the violins represent the posterior distribution of the population trait value and the bars are the median. Each black point is the median trait value for one of the individuals in the base population. Connecting letters above violins indicate populations that significantly different or not. The 95% confidence interval for difference between populations includes 0 for those with the same letter. a. The northernmost populations germinate days later than southern populations in the same greenhouse environment. b. In both South (solid linetype around violin) and North (dashed linetype around violin) gardens, source populations originating from the south had higher probability of winter survival than northern source populations. However, the over-all survival was greater in the South garden, such that even the northernmost source population (Rock Creek) had higher survival in the South than in the North garden.

Fewer plants survived over winter in the North garden than in the South garden (65.5% versus 94.2%). In both gardens, southern source populations survived more often than northern source populations even after accounting for block effects (Fig. 1b). For example, in the North garden, we estimated that plants originating from the most southern source population, Sweetwater River, survived 80.5% (95% CI [71.5%, 88.0%]) of the time compared to 52.3% (95% CI [40.1%, 65.8%]) for the most northern source population Rock Creek. Similarly, in the South garden the local population from Sweetwater River survived 95.2% (95% CI [91.1%, 97.2%]) of the time compared to 85.5% (95% CI [76.4%, 90.8%]) for Rock Creek. For most populations, the average temperature was warmer than the historical normal from 1961–1990, especially for the winter season in the South garden (Fig. S3). However, temperatures for the North garden were close to their historical norm.

### Variance in germination rate and survival within and between populations

The genetic variation in germination rate within populations of *M. cardinalis* is similar to the genetic differences between them (Fig. 2a). The difference between *V*_pop_ and *V*_*G*_ for the log-mean germination rate is −0.017 (95% CI [−0.118, 0.080]), but the confidence intervals include 0. The environmental variance *V*_*E*_ is greater than *V*_*G*_, resulting in a moderate heritability *H*^2^ (Fig. 2b). Maternal and Block effects contributed little variance (Fig. 2a). In contrast, there is virtually no genetic variance in winter survival (*p*_surv_) in either garden, resulting in heritabilities that are effectively 0 (Fig. 3). There is some variation between populations, but it is substantially less than the unexplained environmental variance in both gardens (Fig. 3). Parameter estimates and confidence intervals for all variance components and heritabilities are given in Table S1 for germination rate and Table S2 for winter survival.

**Figure 2:**
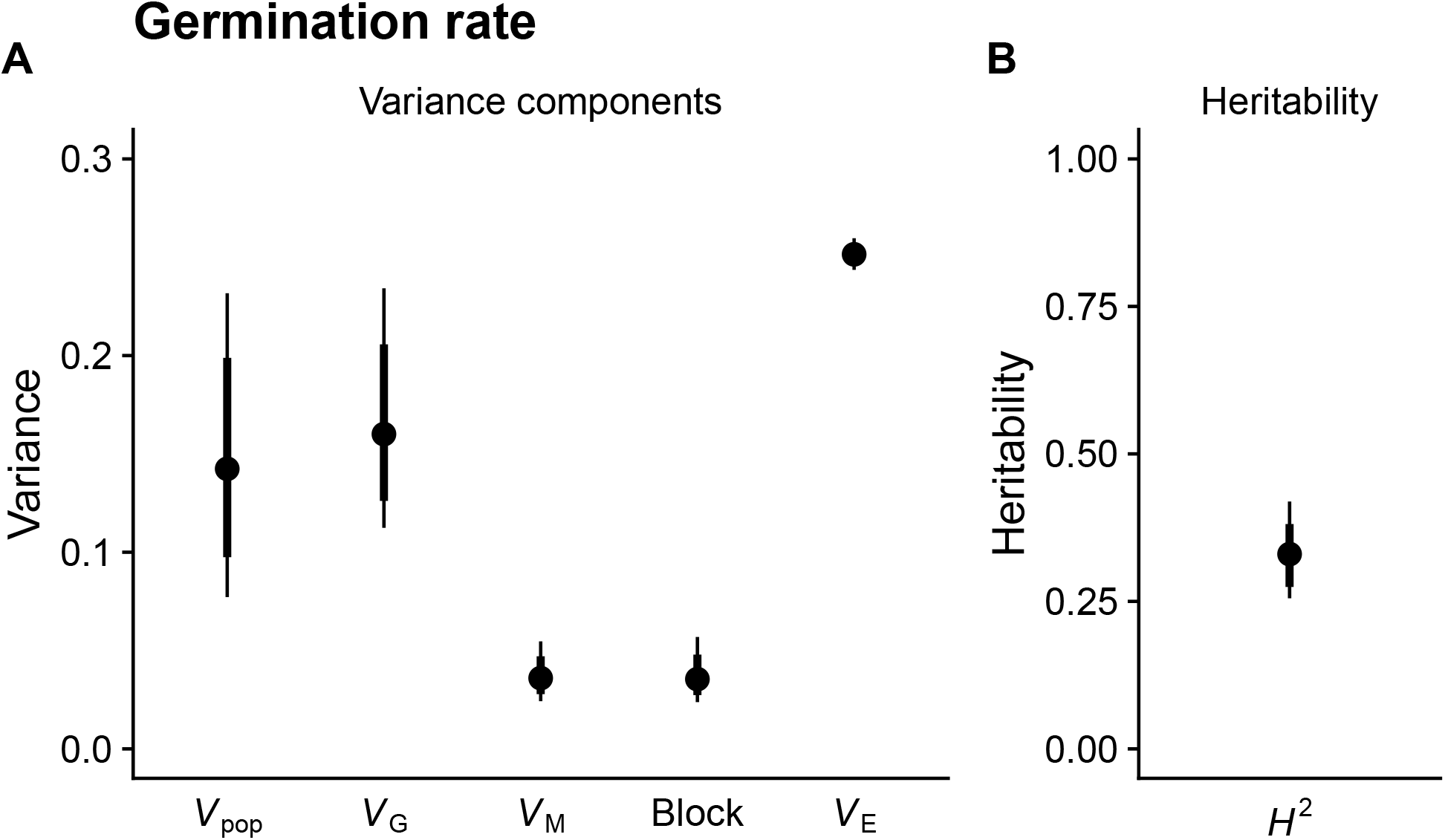
Estimated variance components and heritability of germination rate in units of log(days)^2^. a. The variance in germination rate between source populations *V*_pop_ is comparable to the genetic variation within populations *V*_*G*_. The variance among Blocks and maternal parent (*V*_*M*_) in the greenhouse is substantially lower whereas the unexplained environmental variance is higher *V*_*E*_. b. This results a moderate heritability *H*^2^ which is *V*_*G*_/*V*_*p*_. The point estimates are the median of the posterior distribution; thick lines are 80% confidence intervals; and thin lines are the 95% confidence intervals.

**Figure 3:**
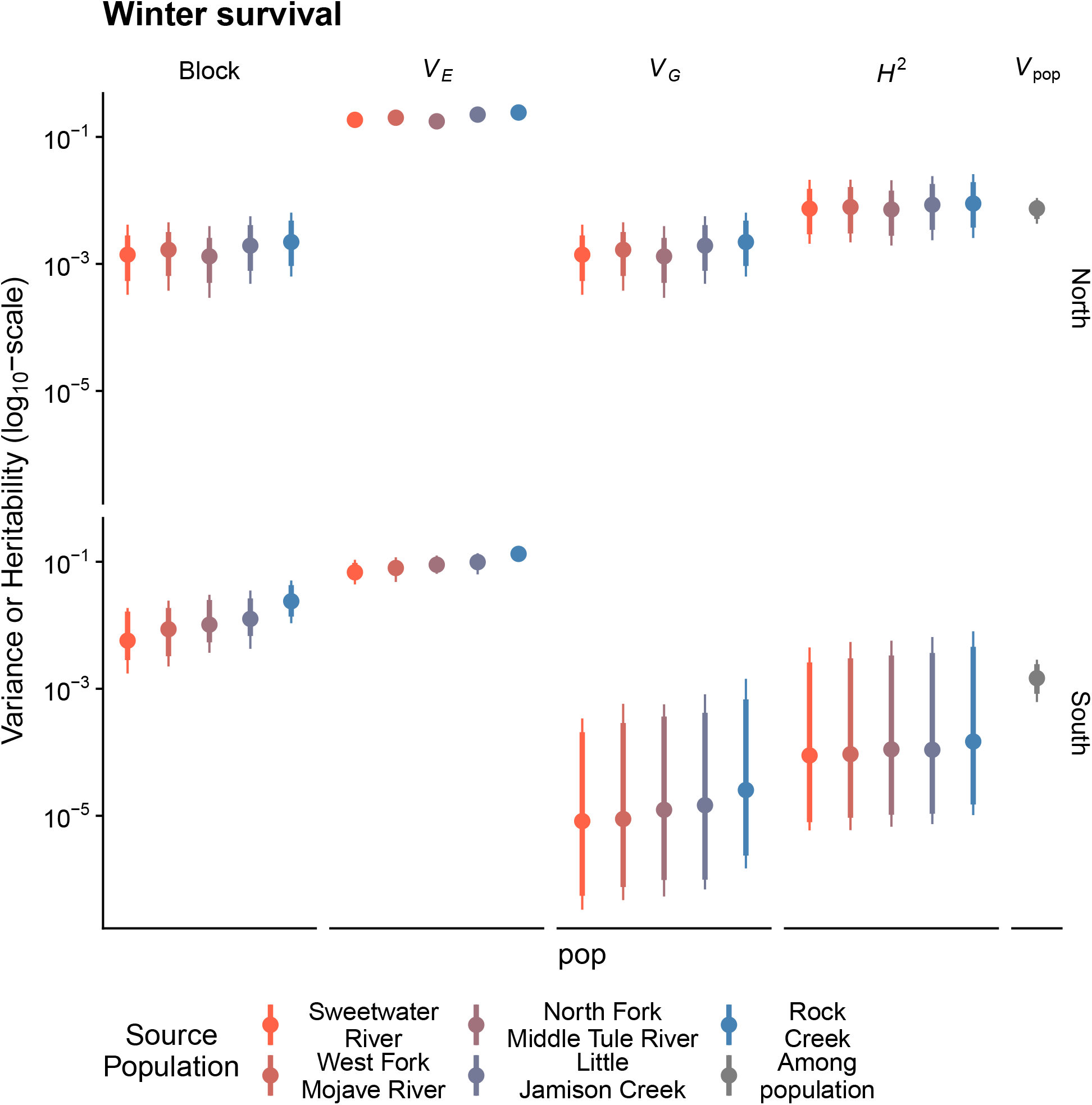
Estimated variance components and heritability of winter survival in units of 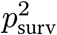. The top row of facets are estimates from the North garden; the bottom row of facets are estimates from the South garden. On the *x*-axis, the source populations are arranged from left (orange) to right (blue) by latitude going from south to north. The right-most facet (*V*_pop_ is grey because it is the variance among populations. On the *y*-axis (log_10_-transformed for visual clarity) is the variance or heritability (*H*^2^) depending on the facet. In both gardens, the unexplained environmental (*V*_*E*_) is higher than the variance contributed by field Block, genetic variance (*V*_*E*_), or variance between source populations (*V*_pop_). Hence, the *H*^2^ is very low in both gardens. The point estimates are the median of the posterior distribution; thick lines are 80% confidence intervals; and thin lines are the 95% confidence intervals.

### Directional selection favors faster germination between but not within populations

Since there was almost no genetic variance in winter survival in either garden, but some variance between populations, the only evidence for selection is between populations. Source populations originating from farther south in the species’ range germinated faster and had higher winter survival than those originating for farther north (Fig. 4). However, within populations, genotypes that germinated faster did not have a different probability of survival. Selection coefficients and confidence intervals are given in Table S3 for between-population selection and Table S4 for genotypic selection. There was no evidence for stabilizing selection using quadratic regression (results not shown).

**Figure 4:**
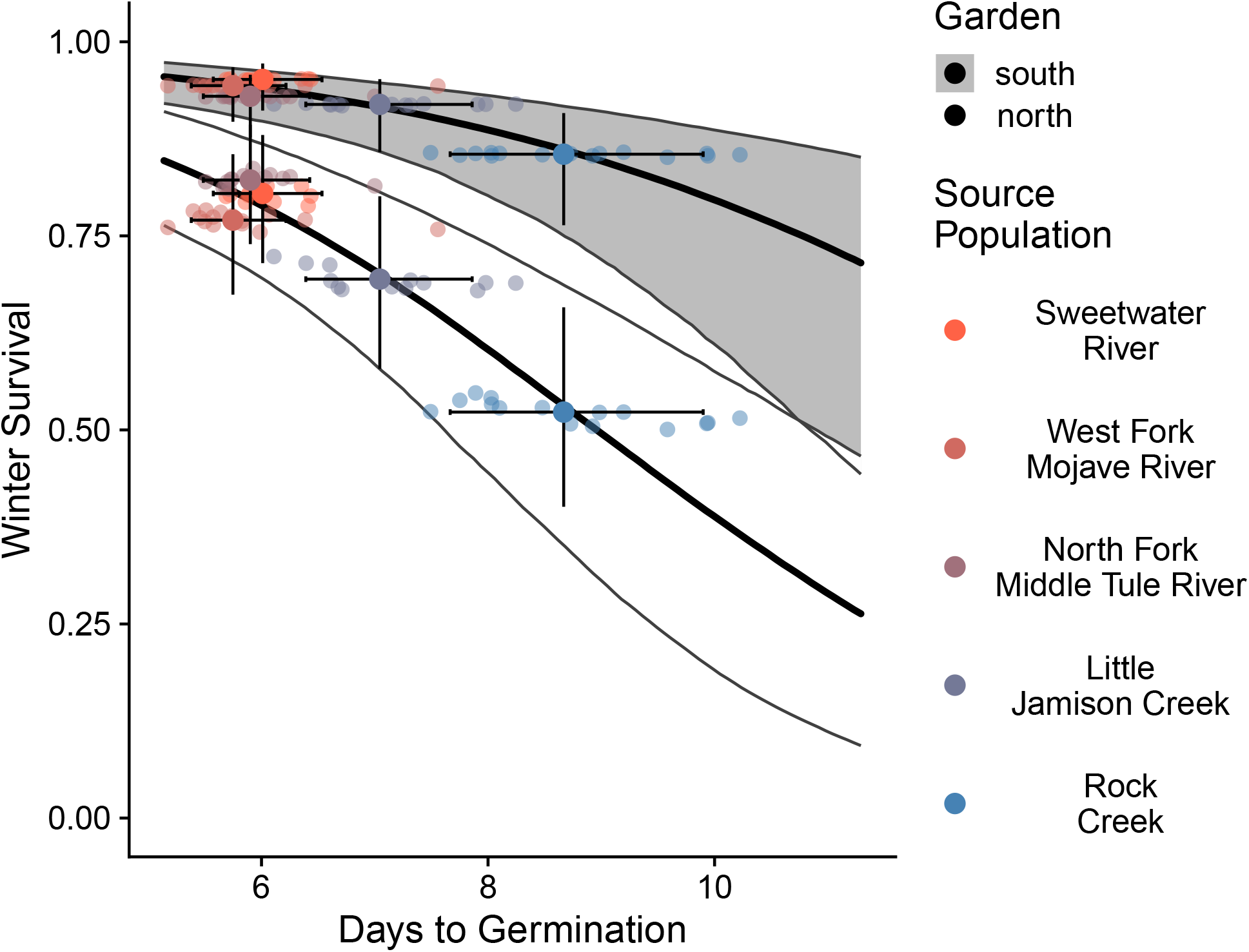
Directional selection favor faster germination between but not within source populations. In both South (grey ribbon) and North (white ribbon) gardens, source populations that germinated faster (*x*-axis) also had higher winter survival (*y*-axis). The larger solid points are the population average estimated from the median of the posterior distribution; smaller translucent points are the genotypic mean trait values within populations. The solid line is regression between germination rate and winter survival between populations estimated from the median of the posterior distribution; the genotypic regression results are not shown in this figure. The ribbon within the thinner black lines is the 95% confidence interval of the regression.

## Discussion

Phenotypic clines are often interpreted as evidence for local adaptation but evolutionary forces besides selection can also generate associations between phenotypes and environment (Endler, 1977). If genotypic selection measured under natural conditions favors local phenotypes, this supports the hypothesis that phenotypic clines are adaptive. Alternatively, failure to detect selection could indicate that nonadaptive evolutionary forces maintain phenotypic variation. We measured the effect of germination rate in the greenhouse, which varies clinally with latitude in *Mimulus cardinalis* (Muir and Angert, 2017), on early winter survival. Although there was genetic variation in germination rate within and between the five source populations (Table 1; Table S1; Fig. 1a), there was no evidence selection consistently favored local genotypes. We predicted that earlier germination should be favored in the South garden and later germination favored in the North garden. Instead, in both North and South gardens, source populations that germinated faster on average also had higher survival, though survival in the North garden was lower than the South overall (Fig. 4). Apparent selection for faster germination in both gardens is likely indirect rather than causal. Populations originating from the southern range limit survived more often and germinated faster. Within populations there was no genetic variance in winter survival despite genetic variance in germination rate, indicating that differences in germination rate are probably not the main cause of differences in winter survival. Our results are not consistent with selection on survival maintaining phenotypic clines in germination rate. Rather, the results may indicate a role for nonadaptive evolution or that selection operates on germination rate during other life stages.

### Why are plants from southern source populations more likely to survive winter?

An unexpected finding is that southern source populations had higher survival in both South and North gardens. If populations were locally adapted, we would expect to see local populations have the highest fitness. We consider three explanations. One explanation is that artificial conditions of the experiment may have favored genotypes from southern populations for an unknown reason. Although we attempted to situate gardens in natural conditions and at realistic conspecific densities, southern genotypes may be better adapted to artificial manipulations such as removing heterospecific competitors, ground cloth, and/or irrigation watering. *M. cardinalis* seeds require moderate temperatures similar to other co-occurring *Mimulus* species (Vickery, 1967) and water, but we do not know the minimum soil water potential required for germination. We also know little about light requirements or soil microbes and experimental coinditions might have differed from natural conditions. A second possibility is that climatic anomalies favor southern, warm-adapted populations in all gardens. Temperature in the North garden was warmer than average for the local population (Little Jamison Creek) but not substantially warmer than the Rock Creek population, which had even lower survival. If we had planted in a season closer to the historical temperature and snowpack, then we might have observed local adaptation in northern source populations. Third, early survival may tradeoff with other fitness components we have not measured or analyzed here, including seedling establishment, later viability, and/or fecundity. We cannot address selection on seedling establishment because we germinated seedlings under controlled greenhouse conditions. In *Arabidopsis thaliana*, faster germination increased survival but decreased seedling establishment (Postma and Ågren, 2018). Southern source population growth rate could be more sensitive to germination timing, leading to stronger response to selection, whereas in the north it matters less and/or conflicting selection at other life stages is stronger. We will address these hypotheses further in a separate paper including multiple seasons of viability and fecundity data from these experiments. We are not including those data in the current analysis because we are using them to address a conceptually distinct question. However, we believe the current analysis advances the field by addressing selection on germination at one life stage and presents a new statistical method for the quantitative genetics of waiting-time traits.

### What maintains genetic variation in germination rate?

One possibility is that germination rate measured in the greenhouse is not representative of what occurs in nature. Species often have particular germination requirements, such as thermal time or water potential (Huang et al., 2016), to escape seed morality, as a bet-hedging strategy in unpredictable environments, or to synchronize establishment with favorable conditions (Donohue et al., 2010). In the field, seeds may use additional cues such as day length or temperature that did not vary in our experiment. We may not be able to measure selection on variation in germination rate unless it occurs in the field. For example, seedlings in the north may require a longer duration of favorable conditions to ensure they do not germinate in late fall or during a brief warm period in early spring. Either scenario might cause them to be exposed to a hard, damaging frost. We may not be able to observe this type of selection without germinating plants in the field at different times of the season. Alternatively, variation may be the result of nonadaptive evolution. *Mimulus cardinalis* demography varies latitudinally in a pattern consistent with an ongoing northward range expansion (Sheth and Angert, 2018). Population expansion can lead to the accumulation of deleterious alleles because of stronger genetic drift (Peischl et al., 2013), serial founder events (Slatkin and Excoffier, 2012), and maladaptation (Polechová, 2018). If selection actually favors faster germination, the slower germination and lower survival of northern populations may both be caused by the accumulation of deleterious genetic variation at the expanding range margins. The current cline could also be a legacy of selection in the recent past which is now less strong or at least not observed during our study (Saccheri et al., 2008). For example, snow addition resembling historical climate increases local advantage in germination and establishment of *Boechera stricta* (Anderson and Wadgymar, 2020). If warming climate is advancing the germination window for *M. cardinalis*, especially in northern Sierra Nevada populations limited by winter snowmelt, this have eliminated any advantage of slower germination rate.

### Statistical approaches for non-Gaussian traits like germination time

Statistical challenges of analyzing non-Gaussian traits may direct research away from “difficult” traits, while applying Gaussian methods to non-Gaussian traits may lead to poor estimation. We show that the discretized log-normal distribution may be useful for analyzing genetic variation in germination time and other “waiting-time” traits with means close to zero. Discretized models take into account that a continuous process is measured at discrete time points (e.g. daily, weekly, etc.). For example, when we saw a seedling had not emerged on day *t* but had emerged by day *t* + 1, our estimation procedure takes into account that germination could have occurred at any time within that interval. Discretization becomes less important when the observation interval is small relative to the total duration, in which case a continuous approximation is sufficient. Waiting-time distributions are naturally bounded at 0 and often skewed, both of which are violated with a Gaussian distribution. We used the log-normal distribution, but other distributions (e.g. exponential or Weibull) might be more appropriate in other situations. A challenge with our approach is that custom probability distributions need to be specified in a programming language. Fortunately, probabilistic programming languages like *Stan* (Stan Development Team, 2022) are enabling evolutionary ecologists and quantitative geneticists to specify custom probability distributions and estimate their parameters from data more easily and reliably (see also Hadfield, 2010).

## Conclusions

The reciprocal transplant experiment did not support our hypothesis that local adaptation maintains clinal variation in germination rate in *M. cardinalis*. We cannot exclude the possibility that artificial aspects of our experimental design may have prevented us from measuring selection on germination rate realistically or that selection operates via different components of fitness. It is also possible that climate anomalies during our experiment prevented us from observing historical patterns of selection that led to present-day phenotypic variation. It is plausible that the cline in germination rate and other phenotypes may be caused by nonadaptive forces that increase the frequency of deleterious alleles during range expansion. Studies like ours that combine clines with reciprocal transplants are necessary to determine what evolutionary forces shape range-wide phenotypic variation within species.

## Acknowledgements

Roger Wynn, Gail Wynn, Christina Schneider, and Norbert Schneider provided land for the gardens. Matthew Bayly, Emily Gercke, Kevin Muir, Rachel Muir, Erin Warkman, and Adam Wilkinson helped with data collection or transplanting. Seema Sheth provided seeds and gave feedback on an earlier draft of the manuscript. Lila Fishman and Kelsey Byers provided suggestions on the non-adaptive cline literature.

## Author Contributions

CDM and ALA designed the experiment. CDM and CLV collected germination data. CDM and ALA collected data on survival in the field. CDM analyzed the data and wrote the manuscript with input from ALA and CLV.

## Data Availability Statement

Data will be deposited on Dryad upon publication. Source code for all analyses will also be archived on Zenodo. Data and code are available to reviewers on GitHub https://github.com/cdmuir/mimulus-germination. The GitHub repo will be revised during revision.

## Supporting Information

Fig. S1-S3 and Tables S1-S4 may be found online in the Supporting Information section.

## Notes

https://github.com/cdmuir/mimulus-germination

### Competing Interest Statement

The authors have declared no competing interest.

