## Supporting Information for "Selection on early survival does not explain germination rate clines in *Mimulus cardinalis* (Phrymaceae)"

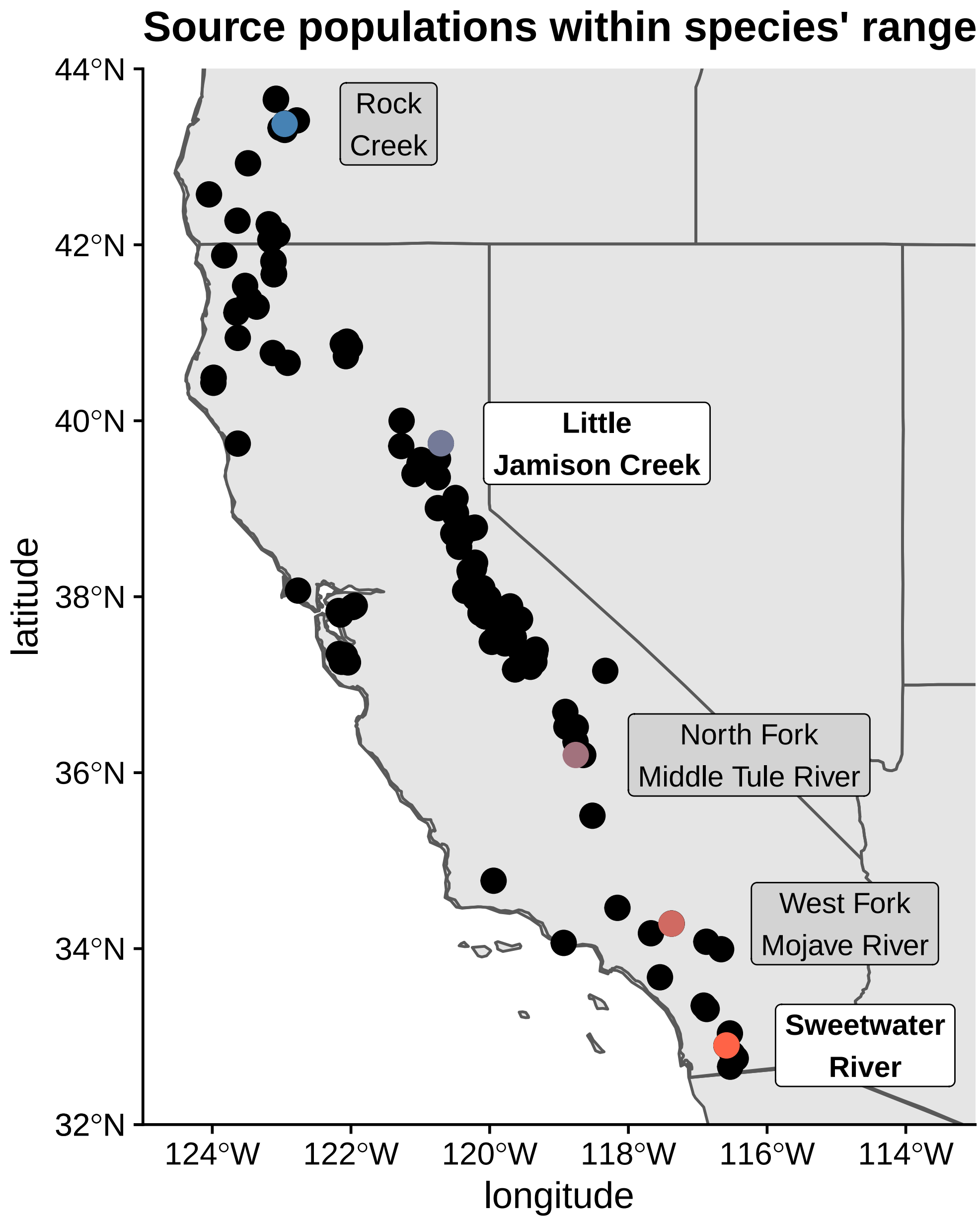


Figure S1: Source populations are distributed through the range of *Mimulus cardinalis*. Dark grey points are occurrence records located throughout California and Oregon, USA (Muir and Angert 2017). Source populations for this study are labeled with colored points. Transplant gardens are labeled with bold font in white boxes.

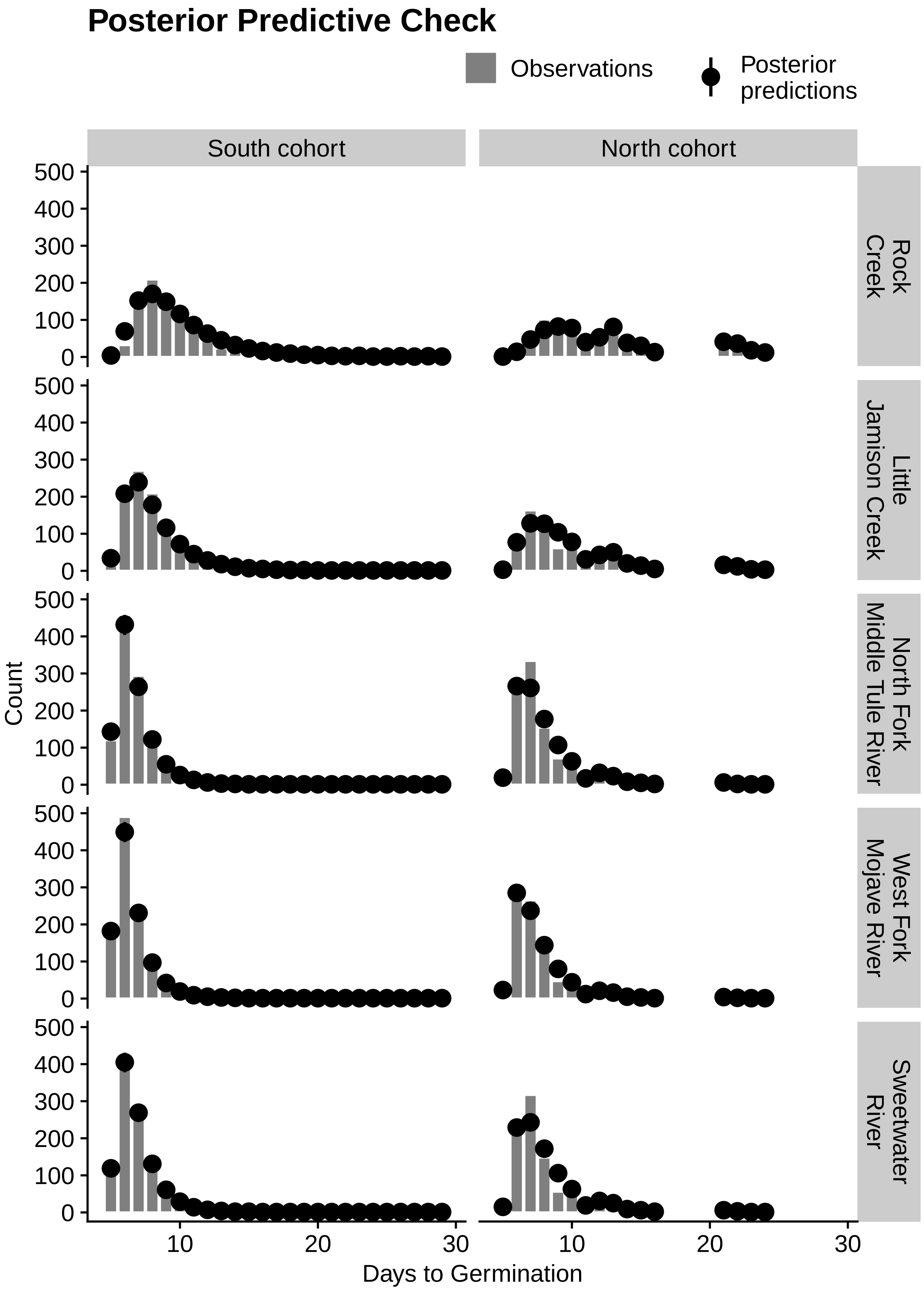


Figure S2: (Caption next page.)

Figure S2: (Previous page.) Posterior predictive check of germination rate from discretized log-normal model for the South cohort (left facets) and North cohort (right facets). Each histogram in gray bars is for one of five *M. cardinalis* populations arranged from northern (top) to southern (bottom). The black points are median values from randomly simulated data sets drawn from each iteration of the posterior distributions. Lines indicate the 95% confidence interval but are usually too narrow to be seen behind the point estimates. A close match between observations and posterior predictions indicate that the model adequately describes the distribution of the data.

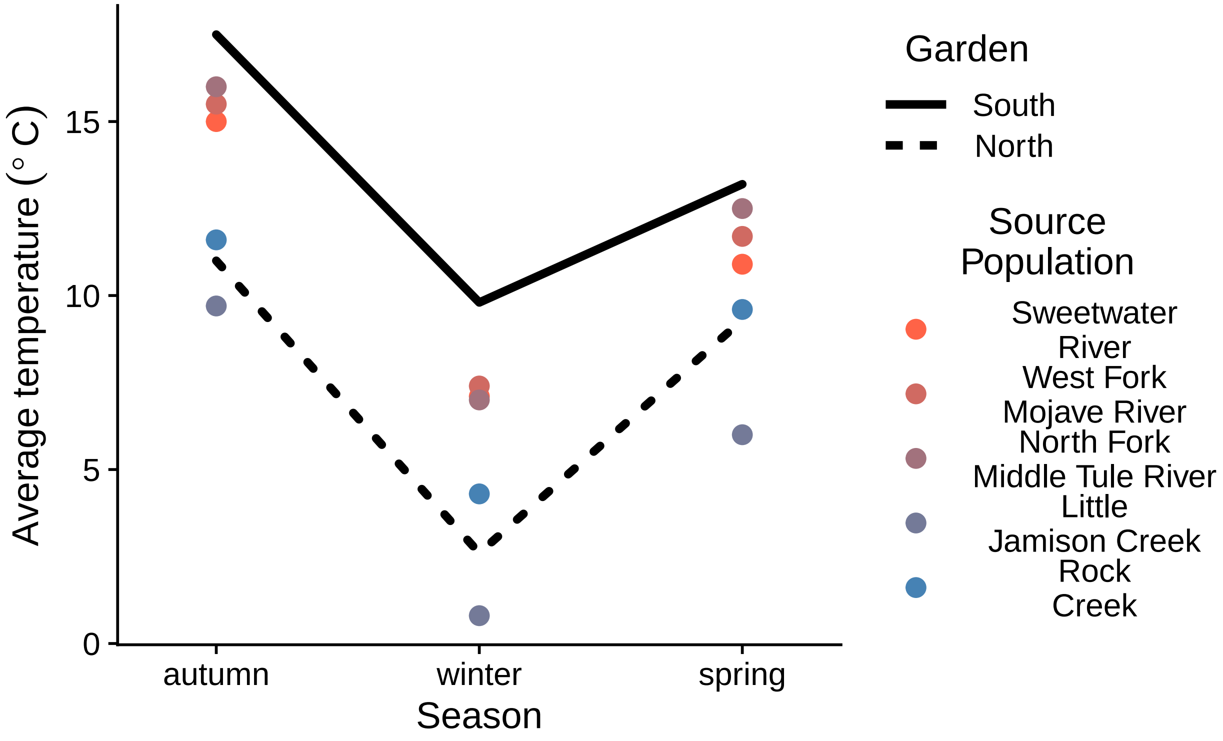


Figure S3: The average temperature during the 2015-16 experiment (lines) was warmer than the historical normal from 1961-1990 (points). The $y$-axis is the average temperature for a season in the southern (solid line) and northern (dashed line) garden. Points are the average seasonal temperature for the 1961-1990 normal for each population source location. Population color is arranged from orange to blue by latitude of origin from south to north. Season is along the $x$-axis. Autumn encompasses September-November; winter encompasses December-February; spring encompasses March-May. All down-scaled climate are from ClimateNA (see Materials and Methods).

Table S1: We decomposed variance in time to germination among populations ($V_{\text{pop}}$), additive genetic differences among individuals in the base population ($V_G$), maternal effects ($V_M$) experimental blocks within the greenhouse (Block), and residual environmental variance ($V_E$). The broad-sense heritability $H^2$ is the proportion of variance contributed by $V_G$. Point estimates and 95% confidence intervals (CI) are estimated from the median and quantiles of the posterior distribution.

| Parameter | Median (95% CI) |
|---|---|
| $V_{\text{pop}}$ | 0.142 (0.0772, 0.232) |
| $V_G$ | 0.16 (0.112, 0.235) |
| $V_M$ | 0.036 (0.0243, 0.0546) |
| Block | 0.0355 (0.0238, 0.0569) |
| $V_E$ | 0.251 (0.244, 0.26) |
| $H^2$ | 0.33 (0.257, 0.422) |

Table S2: We decomposed variance in winter survival (logit($p_{\text{surv}}$)) among populations ($V_{\text{pop}}$), additive genetic differences among individuals in the base population ($V_{\text{G}}$), experimental blocks in the field (Block), and residual environmental variance ($V_{\text{E}}$) for the North and South garden separately. The broad-sense heritability $H^2$ is the proportion of variance contributed by $V_G$. Point estimates and 95% confidence intervals (CI) are estimated from the median and quantiles of the posterior distribution.

| Population | Garden | Parameter | Median (95% CI) |
|---|---|---|---|
| Sweetwater River | North | $H^2$ | $0.00742\ (0.00208,\ 0.0211)$ |
| Sweetwater River | North | $V_{\text{pop}}$ | $0.00744\ (0.00429,\ 0.0109)$ |
| Sweetwater River | North | $V_E$ | $0.186\ (0.144,\ 0.219)$ |
| Sweetwater River | North | $V_G$ | $0.0014\ (0.000327,\ 0.00415)$ |
| Sweetwater River | North | Block | $0.0014\ (0.000327,\ 0.00415)$ |
| West Fork Mojave River | North | $H^2$ | $0.00787\ (0.00219,\ 0.0212)$ |
| West Fork Mojave River | North | $V_{\text{pop}}$ | $0.00744\ (0.00429,\ 0.0109)$ |
| West Fork Mojave River | North | $V_E$ | $0.201\ (0.16,\ 0.228)$ |
| West Fork Mojave River | North | $V_G$ | $0.00167\ (0.000379,\ 0.00453)$ |
| West Fork Mojave River | North | Block | $0.00167\ (0.000379,\ 0.00453)$ |
| North Fork Middle Tule River | North | $H^2$ | $0.0072\ (0.00195,\ 0.0208)$ |
| North Fork Middle Tule River | North | $V_{\text{pop}}$ | $0.00744\ (0.00429,\ 0.0109)$ |
| North Fork Middle Tule River | North | $V_E$ | $0.176\ (0.132,\ 0.212)$ |
| North Fork Middle Tule River | North | $V_G$ | $0.00132\ (0.000292,\ 0.00394)$ |
| North Fork Middle Tule River | North | Block | $0.00132\ (0.000292,\ 0.00394)$ |
| Little Jamison Creek | North | $H^2$ | $0.00854\ (0.00236,\ 0.0241)$ |
| Little Jamison Creek | North | $V_{\text{pop}}$ | $0.00744\ (0.00429,\ 0.0109)$ |
| Little Jamison Creek | North | $V_E$ | $0.224\ (0.185,\ 0.241)$ |
| Little Jamison Creek | North | $V_G$ | $0.00194\ (0.000487,\ 0.00563)$ |
| Little Jamison Creek | North | Block | $0.00194\ (0.000487,\ 0.00563)$ |
| Rock Creek | North | $H^2$ | $0.00894\ (0.00256,\ 0.0259)$ |
| Rock Creek | North | $V_{\text{pop}}$ | $0.00744\ (0.00429,\ 0.0109)$ |
| Rock Creek | North | $V_E$ | $0.242\ (0.228,\ 0.248)$ |
| Rock Creek | North | $V_G$ | $0.00221\ (0.000632,\ 0.00642)$ |
| Rock Creek | North | Block | $0.00221\ (0.000632,\ 0.00642)$ |
| Sweetwater River | South | $H^2$ | $8.91 \times 10^{-5}\ (5.89 \times 10^{-6},\ 0.00448)$ |

| Population | Garden | Parameter | Median (95% CI) |
|---|---|---|---|
| Sweetwater River | South | $V_{\text{pop}}$ | $0.00147\ (0.000618,\ 0.00287)$ |
| Sweetwater River | South | $V_E$ | $0.0683\ (0.044,\ 0.107)$ |
| Sweetwater River | South | $V_G$ | $8.24 \times 10^{-6}\ (3.32 \times 10^{-7},\ 0.000341)$ |
| Sweetwater River | South | Block | $0.00573\ (0.00174,\ 0.0187)$ |
| West Fork Mojave River | South | $H^2$ | $9.34 \times 10^{-5}\ (5.95 \times 10^{-6},\ 0.00545)$ |
| West Fork Mojave River | South | $V_{\text{pop}}$ | $0.00147\ (0.000618,\ 0.00287)$ |
| West Fork Mojave River | South | $V_E$ | $0.0798\ (0.0481,\ 0.117)$ |
| West Fork Mojave River | South | $V_G$ | $8.89 \times 10^{-6}\ (4.7e-07,\ 0.000579)$ |
| West Fork Mojave River | South | Block | $0.00867\ (0.00225,\ 0.0244)$ |
| North Fork Middle Tule River | South | $H^2$ | $0.000111\ (6.73 \times 10^{-6},\ 0.00571)$ |
| North Fork Middle Tule River | South | $V_{\text{pop}}$ | $0.00147\ (0.000618,\ 0.00287)$ |
| North Fork Middle Tule River | South | $V_E$ | $0.0898\ (0.0645,\ 0.125)$ |
| North Fork Middle Tule River | South | $V_G$ | $1.24 \times 10^{-5}\ (5.37 \times 10^{-7},\ 0.000569)$ |
| North Fork Middle Tule River | South | Block | $0.0102\ (0.00369,\ 0.0302)$ |
| Little Jamison Creek | South | $H^2$ | $0.000109\ (7.4e-06,\ 0.00653)$ |
| Little Jamison Creek | South | $V_{\text{pop}}$ | $0.00147\ (0.000618,\ 0.00287)$ |
| Little Jamison Creek | South | $V_E$ | $0.0988\ (0.0633,\ 0.136)$ |
| Little Jamison Creek | South | $V_G$ | $1.47 \times 10^{-5}\ (6.92 \times 10^{-7},\ 0.000819)$ |
| Little Jamison Creek | South | Block | $0.0126\ (0.00426,\ 0.0352)$ |
| Rock Creek | South | $H^2$ | $0.000148\ (1.03 \times 10^{-5},\ 0.00803)$ |
| Rock Creek | South | $V_{\text{pop}}$ | $0.00147\ (0.000618,\ 0.00287)$ |
| Rock Creek | South | $V_E$ | $0.133\ (0.103,\ 0.161)$ |
| Rock Creek | South | $V_G$ | $2.54 \times 10^{-5}\ (1.49 \times 10^{-6},\ 0.00144)$ |
| Rock Creek | South | Block | $0.0239\ (0.0108,\ 0.0507)$ |

Table S3: Between-population directional selection coefficient estimates and confidence intervals. In both North and South gardens, we estimated selection on time to germination (log-mean scale) caused by increased winter survival (logit($p_{\mathrm{surv}}$)). The parameters are the linear regression slope and intercept. Point estimates and 95% confidence intervals (CI) are estimated from the median and quantiles of the posterior distribution.

| Garden | Parameter | Median (95% CI) |
|---|---|---|
| North | intercept | 4.01 (2.97, 5.51) |
| North | slope | −0.444 (−0.675, −0.304) |
| South | intercept | 4.82 (3.73, 6.24) |
| South | slope | −0.344 (−0.556, −0.208) |

Table S4: Genotypic directional selection coefficient estimates and confidence intervals. In both North and South gardens, we estimated selection on time to germination (log-mean scale) caused by increased winter survival (logit($p_{\mathrm{surv}}$)). The parameters are the linear regression slope and intercept. Point estimates and 95% confidence intervals (CI) are estimated from the median and quantiles of the posterior distribution.

| Garden | Parameter | Median (95% CI) |
|---|---|---|
| North | intercept | −0.00151 (−0.0345, 0.0255) |
| North | slope | −0.08 (−0.27, 0.0243) |
| South | intercept | 0.00059 (−0.0156, 0.0244) |
| South | slope | −0.000876 (−0.11, 0.1) |
